# Bacterial Supplements Attenuate Pelvic Irradiation-Induced Brain Metabolic Disruptions via the Gut-Brain Axis: A Multi-Omics Investigation

**DOI:** 10.1101/2024.10.02.616111

**Authors:** Babu Santhi Venkidesh, Rekha Koravadi Narasimhamurthy, Chigateri M. Vinay, Thokur Sreepathy Murali, Kamalesh Dattaram Mumbrekar

## Abstract

Recent advancements in cancer treatments have increased patient survival rates but also led to treatment-related side effects, negatively impacting the quality of life for cancer survivors. Research has highlighted the crucial role of gut microbiota in overall health, including cognition and neurodegenerative disorders. Cancer patients receiving pelvic radiation often experience gut dysbiosis and this may induce changes in brain through the bi-directional connection between the gut microbiota and the brain, known as the microbiota-gut-brain axis. Bacterial supplements intended to enhance health, whether consumed orally or applied topically. However, the mechanism of bacterial supplements to mitigate pelvic radiation-induced metabolomic alterations is not understood. To investigate this, we employed a multi-omics approach to elucidate how these supplements might mitigate radiation-induced metabolomic changes in the rat brain. A single 6 Gy dose of pelvic radiation was administered to 3–4-month-old Sprague Dawley rats and formulated bacterial supplements were given accordingly. Faecal bacterial sequencing and brain metabolomics performed to identify the changes in the gut microbiota and brain metabolomic analysis to check the altered brain metabolites post pelvic radiation. High-throughput 16S rRNA sequencing revealed significant shifts in bacterial composition, with reduced diversity in the radiation group compared to controls, which was restored in the supplementation groups. Notably, the dominant genera in the radiation group included *Methanobrevibacter*, while *Parasutterella* and *Brachyspira* were prevalent in the supplementation cohorts. Untargeted metabolomic analysis identified 2,554 annotated metabolites, with 56 showing significant differences across groups. Principal Component Analysis demonstrated distinct metabolomic profiles between irradiated and control groups, with specific metabolomic pathways like retinol and glycerophospholipid metabolism altered by irradiation. Bacterial supplementation significantly attenuated these metabolomic disruptions. Therefore, bacterial supplementation could be a promising approach to addressing radiation-induced metabolomic reprogramming in the brains through gut dysbiosis in patients undergoing pelvic radiotherapy, enhancing overall well-being.

## Introduction

The intestinal tract is the largest microecosystem in the human body, housing approximately 10^14^ bacteria from over 2,000 known species. These microorganisms collectively contain more than 100 times the genomic DNA of humans (Thursby et al., 2017). These trillions of intestinal inhabitants establish a crucial, evolutionarily driven relationship with the host to maintain homeostasis. Recently, the gut microbiota has gained significant attention in research, particularly its association with various conditions such as cognition and mental health disorders including neurodegenerative disorders (Xiong et al., 2023; Tiwari et al., 2024). The available evidence elucidates bi-directional communication between the gut microbiota and the brain, through the central nervous system, gastrointestinal system, and immune system (Liu et al., 2022). The gut microbiota generates a range of metabolites that supply energy to both itself and the host. These metabolites can influence the host immune and neural functions. Conversely, the gut microbiota is also subject to regulation by the brain functions and metabolomic activities of the host (Chen et al., 2024). This intricate interaction can be disrupted by various factors, including medical treatments.

Pelvic radiotherapy is used to treat cancers located in the pelvic area, such as genitourinary, gynecologic, anorectal, and localized prostate cancers. It is well-known that radiation can cause substantial alterations in the microbial composition of the gut (Fernandes et al., 2023, Joseph et al., 2020). Peliv irradiation can impact gut-brain axis, resulting in a variety of manifestations including neurological consequences, like neuronal death-related brain changes, neuroinflammation and reduced locomotor activity (Venkidesh et al., 2023).

Bacterial supplements have demonstrated the ability to enhance cognitive function and alleviate depressive disorders in both human and animal models by influencing the gut-brain axis (Steenbergen et al., 2015; Ait-Belgnaoui et al., 2014; Venkidesh et al., 2023). However, the complete metabolomic machinery of pelvic radiation-induced neurological manifestation is unknown, and it is necessary to comprehend the role of these bacterial supplements in reversing the metabolomic alterations in the brain to identify novel therapeutic targets and develop treatment techniques, with the potential for personalized medicine on individual metabolic profiles.

We hypothesize that radiation treatment, by depleting the gut microbiome, triggers neuroinflammation in the brain through altered metabolomics which can be mitigated by bacterial supplementation. To accomplish this, the selected bacterial strains were administered orally to pelvic irradiated rats, and the ameliorative effects on the brain metabolome was assessed. Further validation could facilitate the administration of these bacterial strains to individuals undergoing abdominal/pelvic radiotherapy for cancer treatment, who may experience a significant decline in their quality of life.

## Materials and Methods

### Animals

The ethical approval for the utilization of rats for this research was granted by the Institutional Animal Ethics Committee (IAEC/KMC/84/2020), Manipal Academy of Higher Education, Manipal. Male Sprague Dawley rats (3-4 weeks old) weighing 200-250g were maintained under controlled environmental conditions (23±2°C), light (12 hrs of the light/dark cycle), humidity (50±5%), and continuous access to feed and water. Animal care was done under the standards set by WHO, Switzerland, and the Indian National Science Academy, New Delhi, India. The rats were categorized into five groups, each comprising six rats: the control group (C) received no treatment; the radiation group (R) underwent pelvic irradiation of 6 Gy on day 10 of 20-day study period; the BS group received continuous bacterial supplementation throughout the study period; the BS+R group received bacterial supplementation throughout the study, with pelvic irradiation administered on day 10; finally, the R+BS group underwent pelvic irradiation on day 10 followed by bacterial supplementation (Figure. 1).

**Figure 1.**
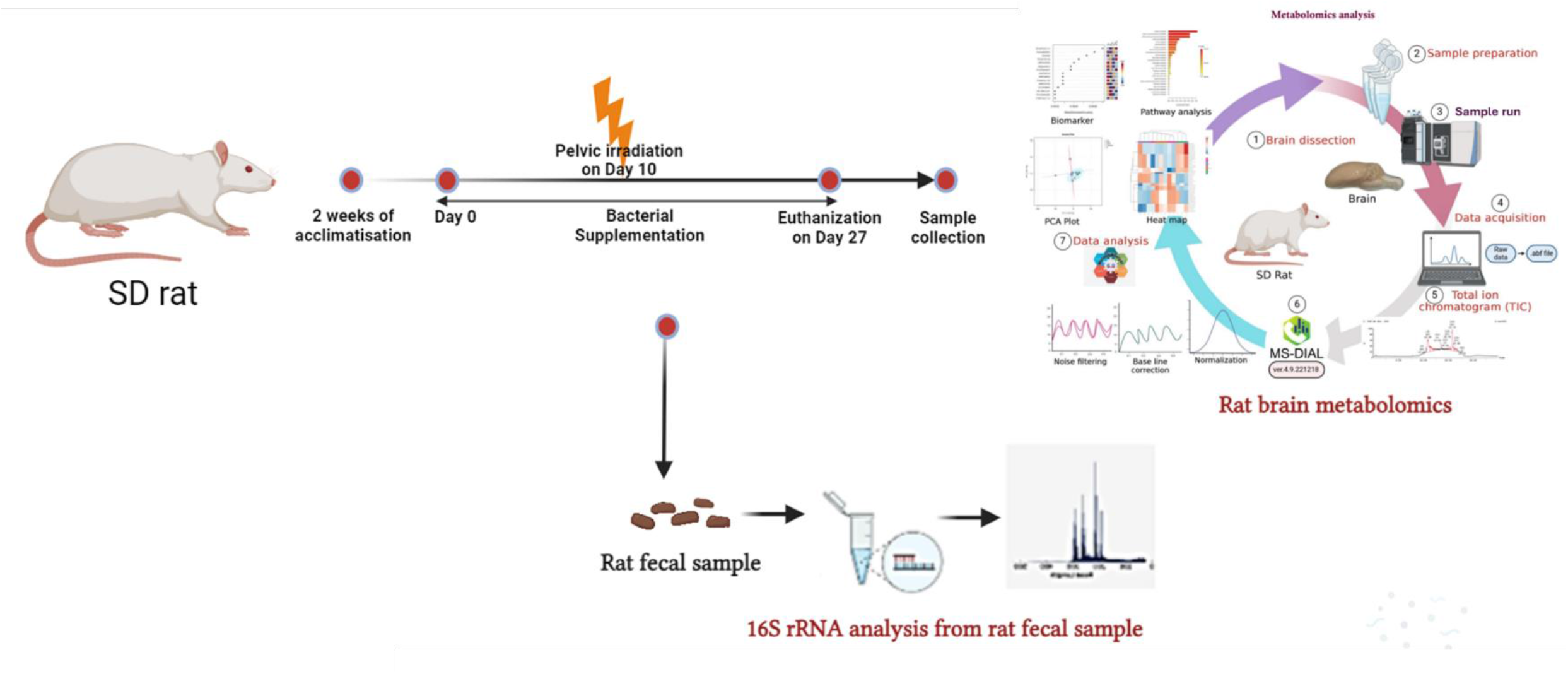
Experimental group and time lines: Sprague Dawley rats were subjected to a single 6 Gy pelvic radiation dose and bacterial supplements were orally dosed. Faecal samples were analyzed for microbial identification, and brain tissues were used for metabolomics analysis.

### Irradiation

Animals in the radiation and combination groups were restrained and exposed to a 6 Gy dose targeting the pelvic region using an Elekta Medical Linear Accelerator (Versa H, Stockholm, Sweden) on the 10^th^ day of the experimental timeline at Shirdi Saibaba Cancer Hospital, Manipal. Additional details regarding the irradiation plan described in Venkidesh et al., 2023.

### Bacterial supplement dosage

A mixed bacterial solution containing *Lactobacillus, Bifidobacterium*, and *Streptococcus*, each at specified CFU levels was orally administered at a dose of 5×10^8^ CFU/mL per day, adjusted according to the rats’ body weight (Venkidesh et al., 2023). Rats in the BS, BS+R, and R+BS groups received these supplements daily throughout the experiment.

### 16S rRNA profiling of rat gut microbiota

Faecal sample collection and DNA isolation were performed according to the previous study (Venkidesh et al., 2023). Further, the sample volume was assessed to ensure they were sufficient for further processing. All samples then underwent a quality check using a spectrophotometer, and the A260/280 ratio was recorded. Additionally, a general PCR targeting the 16S rRNA gene was performed to determine if gene amplification was possible for each sample. Samples that failed the PCR were excluded from further analysis, regardless of passing the spectrophotometric quality control. This step was crucial to ensure the success of the library preparation process. For the PCR successful samples, quantification was conducted using a Qubit fluorometer and a dsDNA broad-range kit. The samples were then diluted to a concentration of 1 ng/µL in a low TE solution. Library preparation was carried out following the Illumina demonstrated protocol for sequencing the V3-V4 region of the 16S rRNA gene. The protocol employed a phased primer strategy based on the approach described by Fadrosh et al. (2014). Indexed libraries were purified and combined at equal concentrations to achieve a concentration of 10nM. The resulting pool of libraries was further validated using an Agilent Tape station 4200 with a high-sensitivity D1000 screen tape and quantified using a Qubit fluorometer with a high-sensitivity dsDNA kit. Subsequently, the pooled library was sequenced on an Illumina MiSeq instrument using the 2×250 base chemistry of the v2 reagent kit, following the manufacturer’s instructions. Demultiplexing of the reads was performed on the sequencer itself, and raw fastq files were generated as the output.

### Raw data processing and taxonomic analysis

Briefly, an initial assessment of raw fastq files was carried out using FastQC v0.11.9 (Andrews et al., 2010). The parameters used to check the raw data quality was provided in a MultiQC report (Ewels et al. 2016) that included information regarding the quantity of data obtained for each of the paired read files for an individual sample. The fastp tool (v0.12.4) (Chen et al. 2018) was used to remove adapter contamination. Each paired-end read was produced using the QIIME pipeline. In order to analyze the species richness, alpha diversity metrices at Observed, Chao-1, Shannon and Simpson were performed. Further analysis will be carried out based on the raw data (Andrews et al., 2010). For processing 16S rRNA gene sequencing data, the initial raw sequence data, referred to as DADA2_input, undergoes several steps to ensure high-quality reads for analysis. This comprehensive process ensured that the final dataset consists of high-quality, accurate, and biologically meaningful sequences, suitable for microbial ecology and diversity studies (Supplementary Table.1). Further, prediction of functional composition of microbial community’s metagenome from its 16S profiles was carried out using PICRUst2 (Douglas et al. 2020) were performed.

### Brain metabolomic sample processing and data analysis

For the brain samples, each 50 mg whole brain sample was mixed with 1,000 μL of 50% cooled methanol solution, and the mixture was homogenized (35 HZ, 60 s, 4°C) and subsequently vortexed (60 s). Then, the sample was placed at -20°C (20 min) to allow the proteins to precipitate. The mixtures were then centrifuged (6,200 ×g, 10 min, 4°C), and all the supernatants transferred to eppendorff tubes were allowed to evaporate until dryness with the stream of nitrogen (35°C). After this, 200 μL of 80% methanol solution was added to the dried residue, and the mixture was completely dissolved and centrifuged (6,200 ×g, 10 min, 4°C), and the supernatant was taken for analysis.

Metabolomic profiling was conducted using the Xevo QTOF ACQUITY UPLC H-Class PLUS Bio System (Waters Corporation, United States) in positive electrospray ionization (ES+) mode. The mobile phase consisted of 0.1% formic acid (FA) in ultrapure water (v/v) for Phase A, and 0.1% FA in acetonitrile (v/v) for Phase B. The positive ionization mode was maintained for 10 minutes at a flow rate of 0.306 mL/min. The gradient mode used was as follows: 0-3.06 minutes: 5% solvent B, 3.06-3.20 minutes: 5-35% solvent B, 3.20-3.54 minutes: 75% solvent B, 3.54-3.67 minutes: 75% solvent B, and 3.67-10 minutes: 5% solvent B. An ACQUITY UPLC BEH C18 column (130 Å, 1.7 μm, 2.1 mm x 50 mm) was employed with an injection volume of 2 μL. The average system pressure was set and maintained at 5093.6 psi.

The obtained LC-MS data were processed in MS Dial 4.9 (Tsugawa et al., 2015) using the following adducts: single charge proton, ammonium, methyl, sodium, potassium, adducts in positive mode using a tolerance limit of ± 15 ppm to improve sensitivity while minimizing background compounds. Using MetaboAnalyst 6.0, PCA plot, heatmap, biomarker and pathway analysis of the altered brain metabolites in all study groups was conducted.

### Statistical analysis

All the data were represented as mean ± SEM. One-way ANOVA with multiple comparison analysis was performed in the GraphPad Prism tool (v9.0, San-Diego, California, USA). A p-value of <0.05 was considered statistically significant.

## Results

### Bacterial supplements reversed pelvic irradiation-induced gut dysbiosis

Illumina 250-bp paired-end sequencing of the amplicon targeting the V3–V4 region of the 16S rRNA gene generated 50k sequencing reads. A quality score greater than or equal to Q30 shows that the base call accuracy at that position is supported by a probability score of 99.9% meaning 1 in 1000 bases at this score can be erroneous (Figure. 2a & 2b). Rats exposed to pelvic irradiation revealed that the mean alpha diversity level of bacterial genera was reduced compared to control and reversed the bacterial composition in the combination group (BS+R) at the Simpson level (Figure. 2c). The distribution of the gut microbiota between the groups was analysed by PCA plot and found that radiation exposure has produced a characteristic dysbiosis compared to the control and BS groups (Figure. 2d). Further, the actual abundance of bacterial genera showed a reduced abundance in *Alloprevotella, Prevotellaceae_group, Bacteroides, Quinella, Eubacterium* in radiation group compared to the control group and reversed after bacterial supplementation (in both BS+R & R+BS) (Figure. 2e). A linear discriminant effect size (LEfSe) analysis showed that different bacterial genera signatures were associated with rats exposed to irradiation compared to that of the rats that received bacterial supplements. The higher prevalence of *Prevotellaceae_UCG_001* was observed in the control group. In the supplementation groups, *Parasutterella, HT002, Streptococcus*, and *Brachyspira* were more prevalent. *Methanobrevibacter* was dominant in the radiation group. In the combination (BS+R) group, *Candidatus_Saccharimonas, Oscillibacter, Parasutterella*, and *Dorea* were observed. *Negativibacillus* was prevalent in the post-irradiation (R+BS) group. Additionally, *Streptococcus and Brachyspira* showed a high linear discriminant analysis (LDA) score greater than 2 (Figure. 2f and 2g). Top 50 MetaCyc pathways per category can be visualized in heatmap showed in Figure. 2h. The top 50 predictions, super pathway of pyrimidine ribonucleotides de novo biosynthesis, super pathway of pyrimidine nucleobases salvage, super pathway of purine nucleotides de novo biosynthesis II, starch degradation V, pentose phosphate pathway (non-oxidative branch), L-lysine biosynthesis III, L-isoleucine biosynthesis IV, gondoate biosynthesis (anaerobic), cis-vaccenate biosynthesis, adenosine ribonucleotides de novo biosynthesis, 5-aminoimidazole ribonucleotide biosynthesis and 5-aminoimidazole ribonucleotide biosynthesis-I were higher in the control groups, however, enrichment of several of these pathways were reduced in the radiation group. Furthermore, the combination group (BS+R) showed a reversal effect on these pathways (Figure 2h).

**Figure 2.**
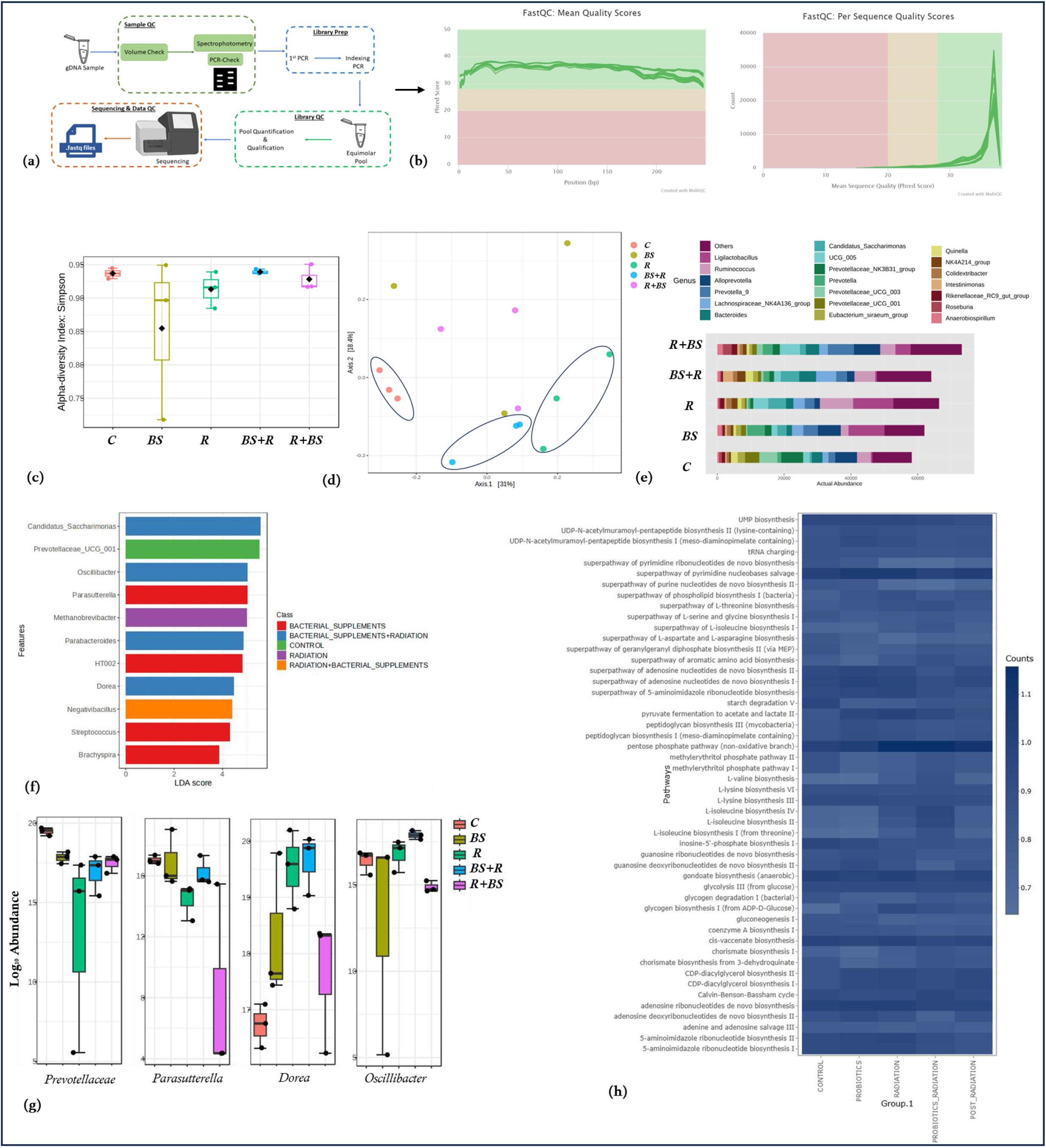
Pelvic irradiation induces gut dysbiosis: (a) Downstream sample processing for the fecal DNA to retrieve the. fastq files, (b) Quality control data showing Phred score and the average counts for the radiation induced gut microbiota and the groups received bacterial supplements, (c) Alpha diversity measure (Simpson) at genus level, (d) Principal Coordinates Analysis (PCoA) plot using the Bray-Curtis distance was conducted to visually examine the similarities and differences in the microbial composition of the samples-p-value: 0.001, (e) Actual microbial abundance after pelvic irradiation and bacterial supplementation, (f) & (g) LEfSe analysis of significant bacterial genera and (h) Prediction of functional composition of microbial community’s metagenome, Top 50 MetaCyc pathways per category can be visualized in heatmap.

### Identification, annotation and ontology of brain metabolites

An untargeted metabolomic analysis was performed on the rat brain across all experimental groups to identify metabolite markers and explore the interaction of microbial metabolites in response to bacterial supplementation in pelvic irradiated rats. LC-MS data of rat brain samples from all the cohorts revealed approximately a total of 7,256 compounds, of which 2,554 were annotated using databases such as HMDB, ChEBI, COCONUT, and FooDB, within a 15ppm range. Among these, 56 compounds were significant across the experimental groups. These metabolites belong to various ontologies, including dialkyldisulfides, piperidines, alkylthiols, dialkylthioethers, ketoximes, N-acyl amines, tertiary carboxylic acids, aldoximes, fatty amides, amides, pyrimidones, triazinones, heteroaromatic compounds, aryl thioethers, hydroxypyrimidines, 1-monoacylglycerols, long-chain fatty acids, long-chain fatty alcohols, 2-monoacylglycerols, 1-alkyl-2-acylglycerols, beta-diketones, organic carbonic acids and derivatives, phenolic glycosides, hybrid peptides, zearalenones, O-cinnamoyl derivatives, macrolactams, diterpene lactones, agarofurans, 1-benzopyrans, triphenyl compounds, phenol ethers, quinolizidines, sesterterpenoids, gamma-amino acids and derivatives, and 1-benzopyrans (Supplementary Table. 2).

### Bacterial supplements attenuated the pelvic irradiation induced rat metabolic alterations in the brain

Principal Component Analysis (PCA) of the metabolomics data revealed a distinct separation between the experimental groups (Fig. 1a). The percent variance explained by the PCA model for Principal Component 1 (PC1) was 20.6%, while Principal Component 2 (PC2) accounted for 15.7% (Fig. 1b). Notably, distinct clusters were observed in the groups exposed to radiation, in contrast to the control and BS group. A heatmap of the top 25 most differentiated brain metabolites showed that 18 of these metabolites were significantly enriched in the irradiation group but were less in the BS combination group (BS+R) (Fig. 1c). Quantitative enrichment analysis (QEA) was performed using metabolite concentrations, employing the global test package version 3 for enrichment analysis. KEGG-based pathway analysis demonstrated higher enrichment of retinol metabolism, glycerophospholipid metabolism, sphingolipid metabolism, steroid hormone biosynthesis and arginine & proline metabolism in between radiation group compared to control. Interestingly, these metabolic pathways remained unaffected in the group that received both bacterial supplementation and radiation (BS+R) (Figure 4) (Supplementary Table. 3 &4).

**Figure 3.**
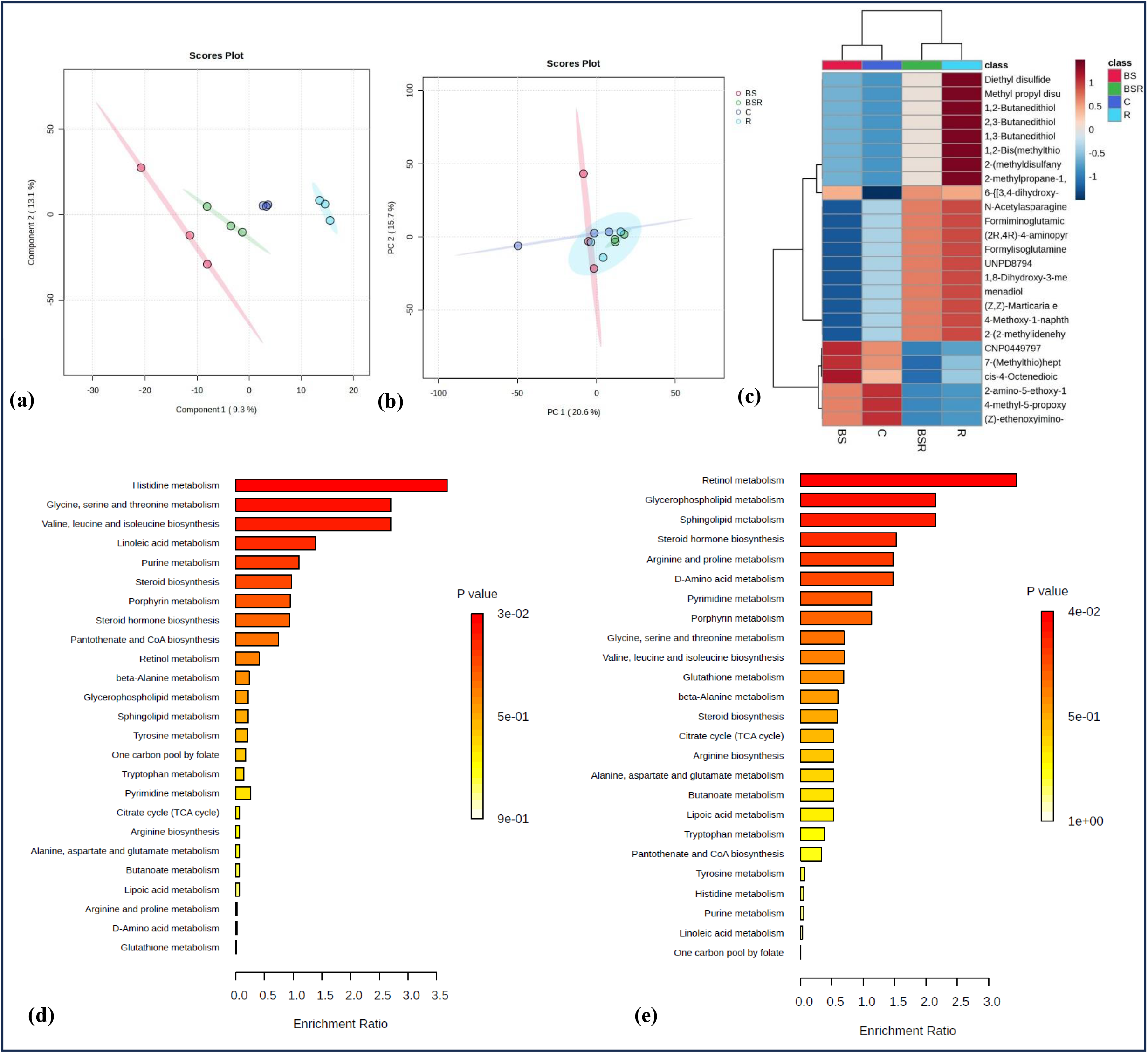
Metabolomic profiling of rat brain using LC/MS analysis after radiation and BS treatment: (a & b) Principal component analysis (PCA) 2D score plot of rat brain metabolites among the groups, (c) Heatmap representing the annotated top 25 metabolite features shows nonsignificant or significant differences in abundance across the treatment groups and (d & e) Enrichment analysis showing significantly enriched pathways of C vs R and R vs BS+R respectively.

**Figure 4.**
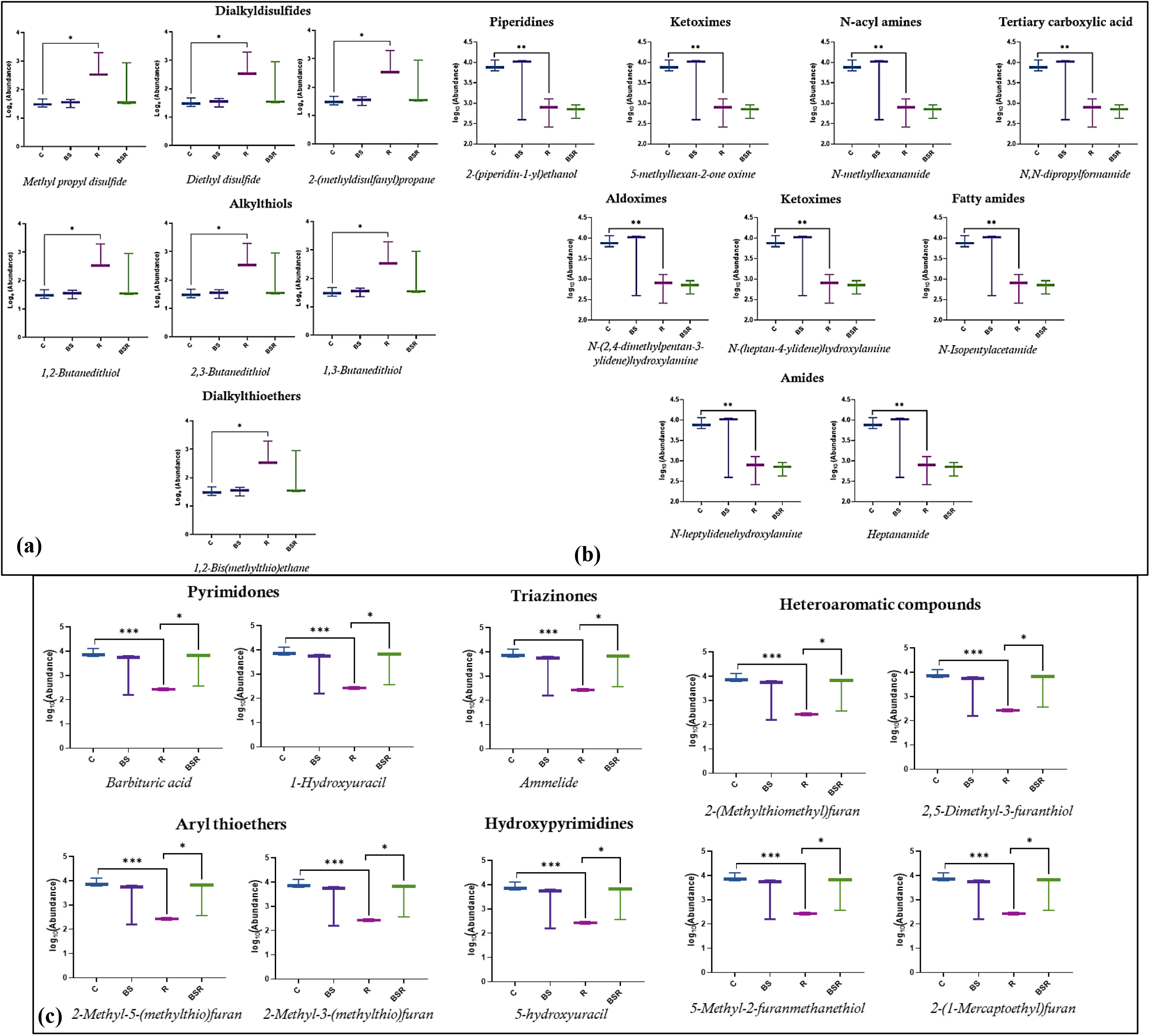
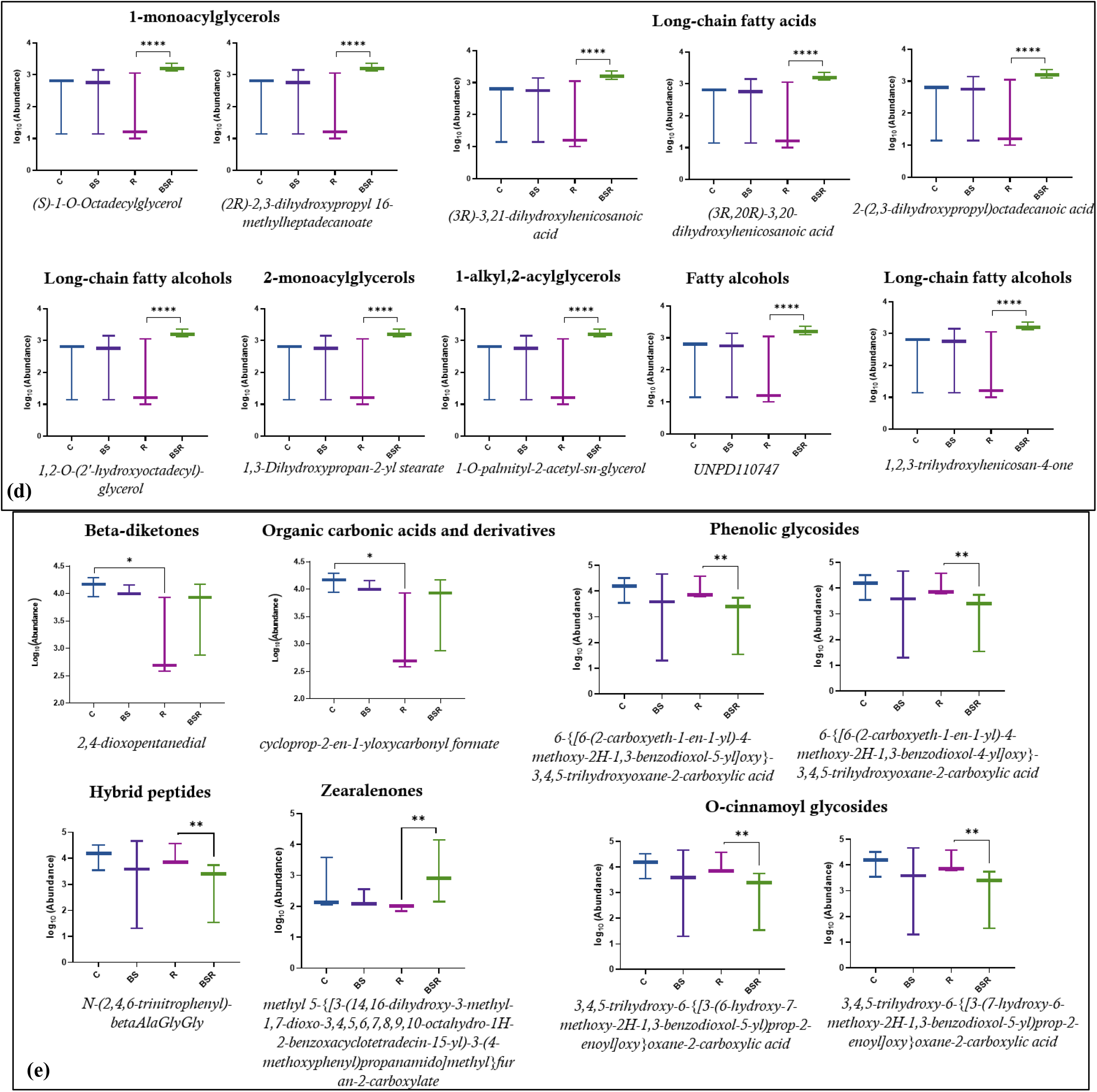
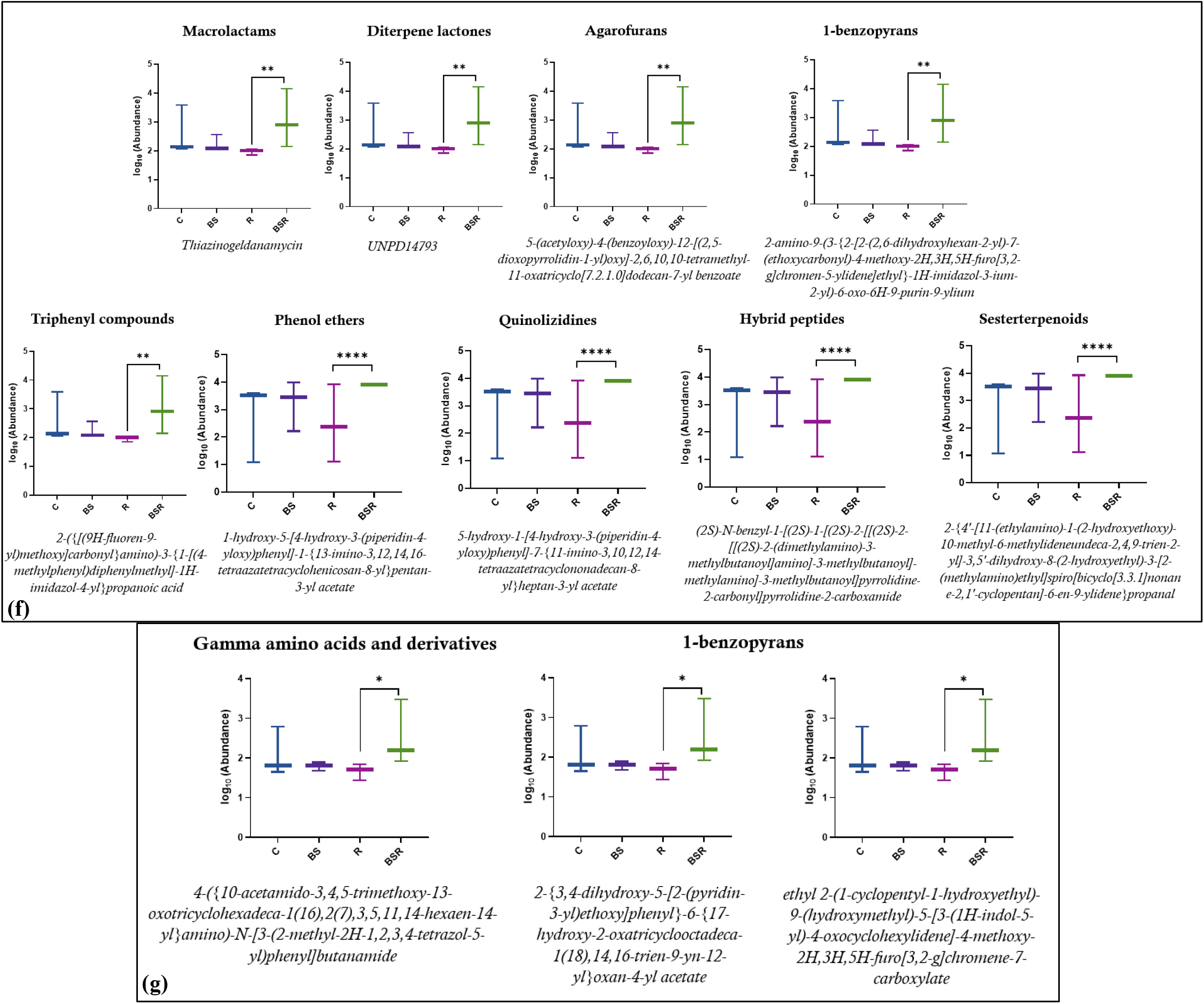
Box-Whisker plot for the significantly differential metabolites in C vs R and R vs BS+R.

The metabolites identified in the irradiated group included various classes such as dialkyldisulfides, alkylthiols, and dialkylthioethers, which were significantly enriched compared to the control group (p< 0.05). Conversely, other metabolite classes, including piperidines, ketoximes, N-acyl amines, tertiary carboxylic acids, aldoximes, fatty amides, and amides, showed lower enrichment in the irradiated group compared to the control (p< 0.05). Upon BS treatment, several brain metabolites, including pyrimidones, triazinones, heteroaromatic compounds, aryl thioethers, and hydroxypyrimidines, were significantly downregulated in the radiation group compared to the control (p < 0.001). However, these metabolites showed a marked improvement in the BS+R group (p < 0.05). Additionally, metabolite classes such as 1-monoacylglycerols, long-chain fatty acids, long-chain fatty alcohols, 2-monoacylglycerols, 1-alkyl-2-acylglycerols, and fatty alcohols were significantly enriched (p < 0.0001). Similarly, phenol ethers, quinolizidines, and sesterterpenoids were also highly enriched (p < 0.0001), alongside gamma-amino acids, their derivatives, and 1-benzopyrans, which were enriched in the BS+R group (p < 0.05) compared to the radiation-only group.

## Discussion

Despite extensive research on the gut-brain axis and its influence on cognition and mental health, the metabolomic pathways involved in radiation-induced neurological damage remain poorly understood. Pelvic radiotherapy is known to disrupt gut microbial composition, leading to neuroinflammation and neuronal damage, yet the specific mechanisms linking these alterations to brain metabolism are unclear. Although bacterial supplementation has shown promise in alleviating cognitive decline and neuroinflammation, there is limited knowledge on how these supplements may reverse the metabolomic changes caused by pelvic radiation. This research gap highlights the need to explore the role of bacterial supplementation in mitigating radiation-induced brain metabolomic alterations, which could lead to novel therapeutic interventions for patients undergoing abdominal or pelvic radiotherapy. In the current study, animals were exposed to pelvic irradiation and further the BS was administered throughout the experiment and in post-irradiation scenario. Gut bacterial changes and importantly the mitigation of brain metabolomic disruptions was studied using genomics and metabolomics approach.

In the current study, the actual abundance of bacterial genera showed a reduced abundance in *Alloprevotella, Prevotellaceae, Bacteroides, Quinella, and Eubacterium* in the radiation group compared to the control group, with a reversal noted after bacterial supplementation (in both BS+R and R+BS). Further LEfSe analysis revealed the control group exhibited a higher prevalence of *Prevotellaceae_UCG_001*, while the supplementation groups showed increased prevalence of *Parasutterella, HT002, Streptococcus, and Brachyspira*. In the radiation group, *Methanobrevibacter* was dominant, whereas the combination (BS+R) group revealed *Candidatus_Saccharimonas, Oscillibacter, Parasutterella, and Dorea. Negativibacillus* was prevalent in the post-irradiation (R+BS) group. Furthermore, *Streptococcus and Brachyspira* demonstrated high LDA scores indicating their significant association with the bacterial supplementation effects. The influence of microbiota on neurological disorders is multifaceted, with specific bacteria playing distinct roles. Studies have shown that *Bifidobacterium* inhibits neuroinflammation by suppressing Aβ accumulation in neurological disorders (Wu et al., 2020). The relative abundance of *Eubacterium* is negatively associated with AD biomarkers, suggesting a protective role (Murray et al., 2022). *Lactobacillus paracasei L9* in Guillain-Barré syndrome down-regulates inflammation-associated metabolites, indicating anti-inflammatory effects (Meng et al., 2023). *Bacteroides fragilis* lipopolysaccharide drives pro-inflammatory pathways linked to neurodegeneration (Lukiw wt al., 2016), while *Akkermansia*’s increased abundance in MS patients correlates with elevated neuroinflammation and low BDNF levels (Lei et al., 2023). Other bacteria, such as *Allobaculum, Alloprevotella, and Bilophila*, have been linked to inflammatory responses and synaptic dysfunction, highlighting the complex interplay between gut microbiota and neurological health (Balaguer-Trias et al., 2022).

In this study, we observed distinct metabolomic features across all the experimental cohorts. The microbial metabolites such as sulfur compounds, furan derivatives, uracils, lipid derivatives, and heterocyclic structures in the brain via the gut-brain axis highlights the intricate role gut microbiota plays in modulating neurological function. Sulfur compounds like methyl propyl disulfide and 1,2-butanedithiol have been shown to affect oxidative stress responses and glutathione regulation, potentially impacting cognitive health (Wang et al., 2022). Similarly, furan derivatives and uracils, produced by gut bacteria during fiber fermentation and nucleic acid metabolism, respectively, can influence neuroinflammatory pathways and DNA repair processes in neurons, offering insight into their role in neurodegenerative diseases. Lipid derivatives like (S)-1-O-Octadecylglycerol are crucial for maintaining neuronal structure and signaling, and their dysregulation has been linked to conditions such as Alzheimer’s disease (Tong et al., 2024). Moreover, antibiotic-like heterocyclic compounds can modulate neurotransmitter pathways, particularly serotonin and dopamine, directly affecting mood and cognition. These findings emphasize the gut microbiome’s significant impact on brain function and the potential therapeutic targets for neurological disorders.

## Conclusion and Future perspectives

Our findings emphasize the importance of bacterial supplements as an important strategy to reverse the pelvic irradiation induced metabolic changes in the brain. Moreover, bacterial supplementation led to a reduction in the presence of enriched metabolites and their associated pathways, indicating that these supplements may serve as neuroprotective agents. Consequently, incorporating bacterial supplements could be a promising strategy for addressing radiation-induced metabolic reprogramming in the brains of patients undergoing pelvic radiotherapy, thereby improving overall well-being. The current study lays the groundwork for future research aimed at improving radiation response outcomes through targeted alterations of the gut microbiome. A molecular docking study could be conducted to investigate the interactions between gut microbial metabolites and proteins associated with neurodegeneration. Further, if any microbial metabolites with positive neuromodulatory or neuroprotective effects are identified, these could be utilized directly to address gut microbiome-induced neurological problems without altering the gut microbiota itself. This approach has the potential to enhance the quality of life for cancer survivors by mitigating neurological issues induced by radiation therapy.

## Author Contributions

Kamalesh Dattaram Mumbrekar & Thokur Sreepathy Murali: designed; conceptualized the study; provided significant comments and revisions to the manuscript. Babu Santhi Venkidesh: maintained the animals; performed and analysed the experiments; and wrote the manuscript. Chigateri M. Vinay: helped in metabolomics analysis, Rekha Koravadi Narasimhamurthy: provided significant contributions to the article.

## Acknowledgments

VBS would like to thank MAHE for the Dr. TMA Pai Ph.D. Fellowship. The authors would like that extend their sincere thanks to Ms. Nenna George, Mrs. Arya, Mr. Sampara Vasishta and Mr. Sanjay K U of MSLS, for their assistance in analysing and troubleshooting the metabolomics data in this study.

## Data availability statement

Not applicable.

## Ethics approval statement

Ethical approval for the study was obtained from the Institutional Animal Ethics Committee (IAEC/KMC/84/2020).

