## Supplementary Table.1, Supplementary Table.2, Supplementary Table.3 & 4, for "Bacterial Supplements Attenuate Pelvic Irradiation-Induced Brain Metabolic Disruptions via the Gut-Brain Axis: A Multi-Omics Investigation": Supplementary table.docx

**Table.1:** **Microbiome profiling by 16S rRNA gene**: First, the reads are filtered to remove low-quality sequences and contaminants, resulting in the filtered data set. Next, the forward (denoisedF) and reverse (denoisedR) reads are denoised to correct sequencing errors and differentiate biological variations, producing amplicon sequence variants (ASVs). These denoised reads are then merged in the merged step, where overlapping regions of the forward and reverse reads are combined to reconstruct the original DNA fragments. Finally, the merged sequences are checked for chimeras, and the nonchim reads, which are non-chimeric, are retained for downstream analysis.

|  | Sample ID | Sample name | DADA2_input | filtered | denoisedF | denoisedR | merged | nonchim |
| --- | --- | --- | --- | --- | --- | --- | --- | --- |
| 1 | M23C203192 | C1 | 54812 | 51968 | 50628 | 48772 | 41519 | 40230 |
| 2 | M23C203193 | C2 | 53243 | 50511 | 49283 | 45813 | 38385 | 36150 |
| 3 | M23C203194 | C3 | 53404 | 50844 | 49333 | 47571 | 39522 | 38135 |
| 4 | M23C203195 | BS1 | 56659 | 29519 | 28441 | 27415 | 23344 | 23202 |
| 5 | M23C203196 | BS2 | 54065 | 45025 | 43849 | 41981 | 34417 | 33904 |
| 6 | M23C203197 | BS3 | 53478 | 51122 | 49708 | 48289 | 39162 | 38856 |
| 7 | M23C203198 | R1 | 49514 | 47297 | 45740 | 45185 | 37465 | 37120 |
| 8 | M23C203199 | R2 | 49414 | 44962 | 43827 | 43659 | 37440 | 37263 |
| 9 | M23C203200 | R3 | 50928 | 45187 | 43730 | 42702 | 34406 | 34115 |
| 10 | M23C203201 | BS+R1 | 51413 | 49115 | 47755 | 47244 | 37486 | 37283 |
| 11 | M23C203202 | BS+R2 | 51151 | 48129 | 46857 | 45942 | 37892 | 37642 |
| 12 | M23C203203 | BS+R3 | 52489 | 49760 | 48292 | 47469 | 38280 | 38168 |
| 13 | M23C203204 | R+BS1 | 53345 | 50122 | 48788 | 47862 | 41269 | 39957 |
| 14 | M23C203205 | R+BS2 | 55751 | 52627 | 51029 | 49514 | 39269 | 39138 |
| 15 | M23C203206 | R+BS3 | 50041 | 47108 | 45766 | 45118 | 37487 | 37300 |

**Table. 2** The compounds identified that are significantly differentially expressed in control and treatment groups

| **Compound Name** | **Ontology** | **Average Rt(min)** | **Average Mz** | **Adduct ion name** | **Databases** | **Formula** |
| --- | --- | --- | --- | --- | --- | --- |
| Methyl propyl disulfide | Dialkyldisulfides | 0.526 | 123.0262 | [M+H]+ | HMDB=HMDB0031872 | C4H10S2 |
| Diethyl disulfide | Dialkyldisulfides | 0.526 | 123.0262 | [M+H]+ | HMDB=HMDB0031872 | C4H10S2 |
| 1,2-Butanedithiol | Alkylthiols | 0.526 | 123.0262 | [M+H]+ | HMDB=HMDB0031872 | C4H10S2 |
| 2,3-Butanedithiol | Alkylthiols | 0.526 | 123.0262 | [M+H]+ | HMDB=HMDB0031872 | C4H10S2 |
| 1,3-Butanedithiol | Alkylthiols | 0.526 | 123.0262 | [M+H]+ | HMDB=HMDB0031872 | C4H10S2 |
| 1,2-Bis(methylthio)ethane | Dialkylthioethers | 0.526 | 123.0262 | [M+H]+ | HMDB=HMDB0031872 | C4H10S2 |
| 2-(methyldisulfanyl)propane | Dialkyldisulfides | 0.526 | 123.0262 | [M+H]+ | HMDB=HMDB0031872 | C4H10S2 |
| Barbituric acid | Pyrimidones | 0.693 | 129.0341 | [M+H]+ | HMDB=HMDB0041833 | C4H4N2O3 |
| 2-Methyl-5-(methylthio)furan | Aryl thioethers | 0.693 | 129.0341 | [M+H]+ | HMDB=HMDB0041833 | C4H4N2O3 |
| ammelide | Triazinones | 0.693 | 129.0341 | [M+H]+ | HMDB=HMDB0041833 | C4H4N2O3 |
| 5-hydroxyuracil | Hydroxypyrimidines | 0.693 | 129.0341 | [M+H]+ | HMDB=HMDB0041833 | C4H4N2O3 |
| 2-(Methylthiomethyl)furan | Heteroaromatic compounds | 0.693 | 129.0341 | [M+H]+ | HMDB=HMDB0041833 | C4H4N2O3 |
| 2,5-Dimethyl-3-furanthiol | Heteroaromatic compounds | 0.693 | 129.0341 | [M+H]+ | HMDB=HMDB0041833 | C4H4N2O3 |
| 1-Hydroxyuracil | Pyrimidones | 0.693 | 129.0341 | [M+H]+ | HMDB=HMDB0041833 | C4H4N2O3 |
| 5-Methyl-2-furanmethanethiol | Heteroaromatic compounds | 0.693 | 129.0341 | [M+H]+ | HMDB=HMDB0041833 | C4H4N2O3 |
| 2-(1-Mercaptoethyl)furan | Heteroaromatic compounds | 0.693 | 129.0341 | [M+H]+ | HMDB=HMDB0041833 | C4H4N2O3 |
| 2-Methyl-3-(methylthio)furan | Aryl thioethers | 0.693 | 129.0341 | [M+H]+ | HMDB=HMDB0041833 | C4H4N2O3 |
| 2-(piperidin-1-yl)ethanol | Piperidines | 0.487 | 130.1279 | [M+H]+ | ChEBI=CHEBI:61238 | C7H15NO |
| 5-methylhexan-2-one oxime | Ketoximes | 0.487 | 130.1279 | [M+H]+ | ChEBI=CHEBI:61238 | C7H15NO |
| N-methylhexanamide | N-acyl amines | 0.487 | 130.1279 | [M+H]+ | ChEBI=CHEBI:61238 | C7H15NO |
| N,N-dipropylformamide | Tertiary carboxylic acid amides | 0.487 | 130.1279 | [M+H]+ | ChEBI=CHEBI:61238 | C7H15NO |
| Heptanamide | Fatty amides | 0.487 | 130.1279 | [M+H]+ | ChEBI=CHEBI:61238 | C7H15NO |
| N-heptylidenehydroxylamine | Aldoximes | 0.487 | 130.1279 | [M+H]+ | ChEBI=CHEBI:61238 | C7H15NO |
| N-(heptan-2-ylidene)hydroxylamine | Ketoximes | 0.487 | 130.1279 | [M+H]+ | ChEBI=CHEBI:61238 | C7H15NO |
| N-(2,4-dimethylpentan-3-ylidene)hydroxylamine | Ketoximes | 0.487 | 130.1279 | [M+H]+ | ChEBI=CHEBI:61238 | C7H15NO |
| N-(heptan-4-ylidene)hydroxylamine | Ketoximes | 0.487 | 130.1279 | [M+H]+ | ChEBI=CHEBI:61238 | C7H15NO |
| N-Isopentylacetamide | Acetamides | 0.487 | 130.1279 | [M+H]+ | ChEBI=CHEBI:61238 | C7H15NO |
| (S)-1-O-Octadecylglycerol | 1-monoacylglycerols |  | 359.3216 | [M+H]+ | HMDB=HMDB00111316052 | C21H42O4 |
| (2R)-2,3-dihydroxypropyl 16-methylheptadecanoate | 1-monoacylglycerols |  | 359.3216 | [M+H]+ | HMDB=HMDB0072842 | C21H42O4 |
| 1,2-O-(2'-hydroxyoctadecyl)-glycerol | Long-chain fatty alcohols | 18.055 | 359.3216 | [M+H]+ | UNPD=UNPD25258 | C21H42O4 |
| 1,3-Dihydroxypropan-2-yl stearate | 2-monoacylglycerols | 18.055 | 359.3216 | [M+H]+ | HMDB=HMDB0011535 | C21H42O4 |
| 1-O-palmityl-2-acetyl-sn-glycerol | 1-alkyl,2-acylglycerols | 18.055 | 359.3216 | [M+H]+ | ChEBI=CHEBI:75936 | C21H42O4 |
| UNPD110747 | Fatty alcohols | 18.055 | 359.3216 | [M+H]+ | UNPD=UNPD110747 | C21H42O4 |
| 1,2,3-trihydroxyhenicosan-4-one | Long-chain fatty alcohols | 18.055 | 359.3216 | [M+H]+ | BLEXP=BLEXPDB00022035687 | C21H42O4 |
| (3R)-3,21-dihydroxyhenicosanoic acid | Long-chain fatty acids | 18.055 | 359.3216 | [M+H]+ | ChEBI=CHEBI:79294 | C21H42O4 |
| (3R,20R)-3,20-dihydroxyhenicosanoic acid | Long-chain fatty acids | 18.055 | 359.3216 | [M+H]+ | ChEBI=CHEBI:79253 | C21H42O4 |
| 2-(2,3-dihydroxypropyl)octadecanoic acid | Long-chain fatty acids | 18.055 | 359.3216 | [M+H]+ | BLEXP=BLEXPDB00009924977 | C21H42O4 |
| 3,4,5-trihydroxy-6-{[3-(6-hydroxy-7-methoxy-2H-1,3-benzodioxol-5-yl)prop-2-enoyl]oxy}oxane-2-carboxylic acid | O-cinnamoyl glycosides | 1.379 | 415.0856 | [M+H]+ | HMDB=HMDB0128726 | C17H18O12 |
| 3,4,5-trihydroxy-6-{[3-(7-hydroxy-6-methoxy-2H-1,3-benzodioxol-5-yl)prop-2-enoyl]oxy}oxane-2-carboxylic acid | O-cinnamoyl glycosides | 1.379 | 415.0856 | [M+H]+ | HMDB=HMDB0128728 | C17H18O12 |
| N-(2,4,6-trinitrophenyl)-betaAlaGlyGly | Hybrid peptides | 1.379 | 415.0856 | [M+H]+ | ChEBI=CHEBI:58980 | C13H14N6O10 |
| 6-{[6-(2-carboxyeth-1-en-1-yl)-4-methoxy-2H-1,3-benzodioxol-5-yl]oxy}-3,4,5-trihydroxyoxane-2-carboxylic acid | Phenolic glycosides | 1.379 | 415.0856 | [M+H]+ | HMDB=HMDB0128725 | C17H18O12 |
| 6-{[6-(2-carboxyeth-1-en-1-yl)-5-methoxy-2H-1,3-benzodioxol-4-yl]oxy}-3,4,5-trihydroxyoxane-2-carboxylic acid | Phenolic glycosides | 1.379 | 415.0856 | [M+H]+ | HMDB=HMDB0128727 | C17H18O12 |
| 3,4,5-trihydroxy-6-{[3-(6-hydroxy-7-methoxy-2H-1,3-benzodioxol-5-yl)prop-2-enoyl]oxy}oxane-2-carboxylic acid | Phenolic glycosides | 1.379 | 415.0856 | [M+H]+ | HMDB=HMDB0128727 | C17H18O12 |
| 3,4,5-trihydroxy-6-{[3-(7-hydroxy-6-methoxy-2H-1,3-benzodioxol-5-yl)prop-2-enoyl]oxy}oxane-2-carboxylic acid | Phenolic glycosides | 1.379 | 415.0856 | [M+H]+ | HMDB=HMDB0128727 | C17H18O12 |
| N-(2,4,6-trinitrophenyl)-betaAlaGlyGly | Phenolic glycosides | 1.379 | 415.0856 | [M+H]+ | HMDB=HMDB0128727 | C17H18O12 |
| 6-{[6-(2-carboxyeth-1-en-1-yl)-4-methoxy-2H-1,3-benzodioxol-5-yl]oxy}-3,4,5-trihydroxyoxane-2-carboxylic acid | Phenolic glycosides | 1.379 | 415.0856 | [M+H]+ | HMDB=HMDB0128727 | C17H18O12 |
| 6-{[6-(2-carboxyeth-1-en-1-yl)-5-methoxy-2H-1,3-benzodioxol-4-yl]oxy}-3,4,5-trihydroxyoxane-2-carboxylic acid | Phenolic glycosides | 1.379 | 415.0856 | [M+H]+ | HMDB=HMDB0128727 | C17H18O12 |
| 2,4-dioxopentanedial | Beta-diketones | 22.353 | 129.0183 | [M+H]+ | BLEXP=BLEXPDB00020591840 | C5H4O4 |
| cycloprop-2-en-1-yloxycarbonyl formate | Organic carbonic acids and derivatives | 22.353 | 129.0183 | [M+H]+ | BLEXP=BLEXPDB00020591840 | C5H4O4 |
| methyl 5-{[3-(14,16-dihydroxy-3-methyl-1,7-dioxo-3,4,5,6,7,8,9,10-octahydro-1H-2-benzoxacyclotetradecin-15-yl)-3-(4-methoxyphenyl)propanamido]methyl}furan-2-carboxylate | Zearalenones | 10.187 | 634.271 | [M+H]+ | COCONUT=CNP0237324 | C35H39NO10 |
| thiazinogeldanamycin | Macrolactams | 10.187 | 634.271 | [M+H]+ | COCONUT=CNP0429460 | C31H43N3O9S |
| UNPD14793 | Diterpene lactones | 10.187 | 634.271 | [M+H]+ | UNPD=UNPD14793,COCONUT=CNP0240969 | C35H39NO10 |
| "5-(acetyloxy)-4-(benzoyloxy)-12-[(2,5-dioxopyrrolidin-1-yl)oxy]-2,6,10,10-tetramethyl-11-oxatricyclo[7.2.1.0]dodecan-7-yl benzoate" | Agarofurans | 10.187 | 634.271 | [M+H]+ | COCONUT=CNP0078786 | C35H39NO10 |
| 2-amino-9-(3-{2-[2-(2,6-dihydroxyhexan-2-yl)-7-(ethoxycarbonyl)-4-methoxy-2H,3H,5H-furo[3,2-g]chromen-5-ylidene]ethyl}-1H-imidazol-3-ium-2-yl)-6-oxo-6H-9-purin-9-ylium | 1-benzopyrans | 10.187 | 634.271 | [M+H]+ | COCONUT=CNP0009789;CNP0256224 | C31H35N7O8 |
| 2-({[(9H-fluoren-9-yl)methoxy]carbonyl}amino)-3-{1-[(4-methylphenyl)diphenylmethyl]-1H-imidazol-4-yl}propanoic acid | Triphenyl compounds | 10.187 | 634.271 | [M+H]+ | COCONUT=CNP0058865 | C41H35N3O4 |
| 1-hydroxy-5-[4-hydroxy-3-(piperidin-4-yloxy)phenyl]-1-{13-imino-3,12,14,16-tetraazatetracyclohenicosan-8-yl}pentan-3-yl acetate | Phenol ethers | 10.187 | 634.271 | [M+H]+ | UNPD=UNPD123863,COCONUT=CNP0307208 | C38H64O7 |
| 5-hydroxy-1-[4-hydroxy-3-(piperidin-4-yloxy)phenyl]-7-{11-imino-3,10,12,14-tetraazatetracyclononadecan-8-yl}heptan-3-yl acetate | Quinolizidines | 10.187 | 634.271 | [M+H]+ | UNPD=UNPD86583,COCONUT=CNP0377283 | C38H64O7 |
| (2S)-N-benzyl-1-[(2S)-1-[(2S)-2-[[(2S)-2-[[(2S)-2-(dimethylamino)-3-methylbutanoyl]amino]-3-methylbutanoyl]-methylamino]-3-methylbutanoyl]pyrrolidine-2-carbonyl]pyrrolidine-2-carboxamide | Hybrid peptides | 10.187 | 634.271 | [M+H]+ | UNPD=UNPD113951,COCONUT=CNP0270260 | C38H64O7 |
| 2-{4'-[11-(ethylamino)-1-(2-hydroxyethoxy)-10-methyl-6-methylideneundeca-2,4,9-trien-2-yl]-3,5'-dihydroxy-8-(2-hydroxyethyl)-3-[2-(methylamino)ethyl]spiro[bicyclo[3.3.1]nonane-2,1'-cyclopentan]-6-en-9-ylidene}propanal | Sesterterpenoids | 10.187 | 634.271 | [M+H]+ | UNPD=UNPD113951,COCONUT=CNP0270260 | C38H64O7 |
| 4-({10-acetamido-3,4,5-trimethoxy-13-oxotricyclohexadeca-1(16),2(7),3,5,11,14-hexaen-14-yl}amino)-N-[3-(2-methyl-2H-1,2,3,4-tetrazol-5-yl)phenyl]butanamide | Gamma amino acids and derivatives | 10.184 | 650.2657 | [M+Na]+ | COCONUT=CNP0408117 | C33H37N7O6 |
| 2-{3,4-dihydroxy-5-[2-(pyridin-3-yl)ethoxy]phenyl}-6-{17-hydroxy-2-oxatricyclooctadeca-1(18),14,16-trien-9-yn-12-yl}oxan-4-yl acetate | Catechols | 10.184 | 650.2657 | [M+Na]+ | COCONUT=CNP0240422 | C37H41NO8 |
| ethyl 2-(1-cyclopentyl-1-hydroxyethyl)-9-(hydroxymethyl)-5-[3-(1H-indol-5-yl)-4-oxocyclohexylidene]-4-methoxy-2H,3H,5H-furo[3,2-g]chromene-7-carboxylate | 1-benzopyrans | 10.184 | 650.2657 | [M+Na]+ | COCONUT=CNP0337322 | C37H41NO8 |

**Table.3:** The table below shows the detailed results from the pathway analysis (C vs R).

|  | **Total compound** | **Hits** | **Raw p** | **-log10**  **(p)** | **Holm adjust** | **FDR** | **Impact** |
| --- | --- | --- | --- | --- | --- | --- | --- |
| Retinol metabolism | 17 | 1 | 69.04 | 20.00 | 4.05E-02 | 1.00E+00 | 7.40E-01 |
| Glycerophospholipid metabolism | 36 | 1 | 43.14 | 20.00 | 1.56E-01 | 1.00E+00 | 7.40E-01 |
| Sphingolipid metabolism | 32 | 1 | 43.14 | 20.00 | 1.56E-01 | 1.00E+00 | 7.40E-01 |
| Steroid hormone biosynthesis | 87 | 5 | 30.60 | 20.00 | 2.47E-01 | 1.00E+00 | 7.40E-01 |
| Arginine and proline metabolism | 36 | 1 | 29.58 | 20.00 | 2.65E-01 | 1.00E+00 | 7.40E-01 |
| D-Amino acid metabolism | 15 | 1 | 29.58 | 20.00 | 2.65E-01 | 1.00E+00 | 7.40E-01 |
| Pyrimidine metabolism | 39 | 2 | 22.60 | 20.00 | 3.38E-01 | 1.00E+00 | 7.40E-01 |
| Porphyrin metabolism | 31 | 1 | 22.55 | 20.00 | 3.41E-01 | 1.00E+00 | 7.40E-01 |
| Glycine, serine and threonine metabolism | 33 | 1 | 13.87 | 20.00 | 4.67E-01 | 1.00E+00 | 7.40E-01 |
| Valine, leucine and isoleucine biosynthesis | 8 | 1 | 13.87 | 20.00 | 4.67E-01 | 1.00E+00 | 7.40E-01 |
| Glutathione metabolism | 28 | 1 | 13.83 | 20.00 | 4.68E-01 | 1.00E+00 | 7.40E-01 |
| beta-Alanine metabolism | 21 | 1 | 11.88 | 20.00 | 5.04E-01 | 1.00E+00 | 7.40E-01 |
| Steroid biosynthesis | 41 | 5 | 11.62 | 20.00 | 5.08E-01 | 1.00E+00 | 7.40E-01 |
| Citrate cycle (TCA cycle) | 20 | 1 | 10.42 | 20.00 | 5.33E-01 | 1.00E+00 | 7.40E-01 |
| Arginine biosynthesis | 14 | 1 | 10.42 | 20.00 | 5.33E-01 | 1.00E+00 | 7.40E-01 |
| Alanine, aspartate and glutamate metabolism | 28 | 1 | 10.42 | 20.00 | 5.33E-01 | 1.00E+00 | 7.40E-01 |
| Butanoate metabolism | 15 | 1 | 10.42 | 20.00 | 5.33E-01 | 1.00E+00 | 7.40E-01 |
| Lipoic acid metabolism | 28 | 1 | 10.42 | 20.00 | 5.33E-01 | 1.00E+00 | 7.40E-01 |
| Tryptophan metabolism | 41 | 1 | 7.59 | 20.00 | 5.97E-01 | 1.00E+00 | 7.40E-01 |
| Pantothenate and CoA biosynthesis | 20 | 2 | 6.69 | 20.00 | 8.06E-01 | 1.00E+00 | 7.40E-01 |
| Tyrosine metabolism | 42 | 1 | 1.26 | 20.00 | 8.32E-01 | 1.00E+00 | 7.40E-01 |
| Histidine metabolism | 16 | 1 | 0.98 | 20.00 | 8.52E-01 | 1.00E+00 | 7.40E-01 |
| Purine metabolism | 70 | 1 | 0.96 | 20.00 | 8.53E-01 | 1.00E+00 | 7.40E-01 |
| Linoleic acid metabolism | 5 | 1 | 0.52 | 20.00 | 8.92E-01 | 1.00E+00 | 7.40E-01 |
| One carbon pool by folate | 9 | 1 | 0.08 | 20.00 | 8.59E-01 | 1.00E+00 | 7.40E-01 |

**Table.4:** The table below shows the detailed results from the pathway analysis (R vs BS+R).

|  | **Total compound** | **Hits** | **Raw p** | **-log10**  **(p)** | **Holm adjust** | **FDR** | **Impact** |
| --- | --- | --- | --- | --- | --- | --- | --- |
| Histidine metabolism | 16 | 1 | 73.72 | 20.00 | 2.86E-02 | 7.14E-01 | 7.14E-01 |
| Glycine, serine and threonine metabolism | 33 | 1 | 54.02 | 20.00 | 9.61E-02 | 1.00E+00 | 8.00E-01 |
| Valine, leucine and isoleucine biosynthesis | 8 | 1 | 54.02 | 20.00 | 9.61E-02 | 1.00E+00 | 8.00E-01 |
| Linoleic acid metabolism | 5 | 1 | 27.94 | 20.00 | 2.81E-01 | 1.00E+00 | 9.29E-01 |
| Purine metabolism | 70 | 1 | 22.05 | 20.00 | 3.47E-01 | 1.00E+00 | 9.29E-01 |
| Steroid biosynthesis | 41 | 5 | 19.43 | 20.00 | 3.82E-01 | 1.00E+00 | 9.29E-01 |
| Porphyrin metabolism | 31 | 1 | 18.93 | 20.00 | 3.89E-01 | 1.00E+00 | 9.29E-01 |
| Steroid hormone biosynthesis | 87 | 5 | 18.84 | 20.00 | 4.11E-01 | 1.00E+00 | 9.29E-01 |
| Pantothenate and CoA biosynthesis | 20 | 2 | 14.99 | 20.00 | 5.57E-01 | 1.00E+00 | 9.29E-01 |
| Retinol metabolism | 17 | 1 | 8.12 | 20.00 | 5.84E-01 | 1.00E+00 | 9.29E-01 |
| beta-Alanine metabolism | 21 | 1 | 4.65 | 20.00 | 6.82E-01 | 1.00E+00 | 9.29E-01 |
| Glycerophospholipid metabolism | 36 | 1 | 4.34 | 20.00 | 6.92E-01 | 1.00E+00 | 9.29E-01 |
| Sphingolipid metabolism | 32 | 1 | 4.34 | 20.00 | 6.92E-01 | 1.00E+00 | 9.29E-01 |
| Tyrosine metabolism | 42 | 1 | 4.17 | 20.00 | 6.98E-01 | 1.00E+00 | 9.29E-01 |
| One carbon pool by folate | 9 | 1 | 3.35 | 20.00 | 7.29E-01 | 1.00E+00 | 9.29E-01 |
| Tryptophan metabolism | 41 | 1 | 2.72 | 20.00 | 7.55E-01 | 1.00E+00 | 9.29E-01 |
| Pyrimidine metabolism | 39 | 2 | 5.21 | 20.00 | 7.76E-01 | 1.00E+00 | 9.29E-01 |
| Citrate cycle (TCA cycle) | 20 | 1 | 1.21 | 20.00 | 8.36E-01 | 1.00E+00 | 9.29E-01 |
| Arginine biosynthesis | 14 | 1 | 1.21 | 20.00 | 8.36E-01 | 1.00E+00 | 9.29E-01 |
| Alanine, aspartate and glutamate metabolism | 28 | 1 | 1.21 | 20.00 | 8.36E-01 | 1.00E+00 | 9.29E-01 |
| Butanoate metabolism | 15 | 1 | 1.21 | 20.00 | 8.36E-01 | 1.00E+00 | 9.29E-01 |
| Lipoic acid metabolism | 28 | 1 | 1.21 | 20.00 | 8.36E-01 | 1.00E+00 | 9.29E-01 |
| Arginine and proline metabolism | 36 | 1 | 0.52 | 20.00 | 8.92E-01 | 1.00E+00 | 9.29E-01 |
| D-Amino acid metabolism | 15 | 1 | 0.52 | 20.00 | 8.92E-01 | 1.00E+00 | 9.29E-01 |
| Glutathione metabolism | 28 | 1 | 0.14 | 20.00 | 9.45E-01 | 1.00E+00 | 9.45E-01 |
